# Hierarchical optimization for the efficient parametrization of ODE models

**DOI:** 10.1101/247924

**Authors:** Carolin Loos, Sabrina Krause, Jan Hasenauer

## Abstract

Mathematical models are nowadays important tools for analyzing dynamics of cellular processes. The unknown model parameters are usually estimated from experimental data. These data often only provide information about the relative changes between conditions, hence, the observables contain scaling parameters. The unknown scaling parameters and corresponding noise parameters have to be inferred along with the dynamic parameters. The nuisance parameters often increase the dimensionality of the estimation problem substantially and cause convergence problems. In this manuscript, we propose a hierarchical optimization approach for estimating the parameters for ordinary differential equation (ODE) models from relative data. Our approach restructures the optimization problem into an inner and outer subproblem. These subproblems possess lower dimensions than the original optimization problem, and the inner problem can be solved analytically. We evaluated accuracy, robustness, and computational efficiency of the hierarchical approach by studying three signaling pathways. The proposed approach achieved better convergence than the standard approach and required a lower computation time. As the hierarchical optimization approach is widely applicable, it provides a powerful alternative to established approaches.

## 1 Introduction

Mechanistic mathematical models are used in systems biology to improve the understanding of biological processes. The mathematical models most frequently used in systems biology are probably ordinary differential equations (ODEs). ODE models are, among others, used to describe the dynamics of biochemical reaction networks (Kitano, 2002; Klipp et al., 2005; Sch¨oberl et al., 2009) and proliferation/differentiation processes (De Boer et al., 2006). The dynamic parameters of the underlying processes, e.g., reaction rates and initial conditions, are often unknown and need to be inferred from available experimental data. The inference provides information about the plausibility of the model topology, and the inferred parameters might for instance be used to predict latent variables or the response of the process to perturbations (Molinelli et al., 2013).

The experimental data used for parameter estimation are produced by various experimental techniques. Most of these techniques provide relative data, meaning that the observation is proportional to a variable of interest, e.g., the concentration of a chemical species. This is for instance the case for Western blotting (Renart et al., 1979) and flow and mass cytometry (Herzenberg et al., 2006). If calibration curves are generated, the measured intensities can be converted to concentrations, however, in most studies this is not done due to increased resource demands.

In the literature, two methods are employed to link relative data to mathematical models: (i) evaluation of relative changes (Degasperi et al., 2017) and (ii) introduction of scaling parameters (Raue et al., 2013). In (i), relative changes between conditions are compared, and the differences between observed and simulated relative changes are minimized. While this approach is intuitive and does not alter the dimension of the fitting problem, the noise distribution is non-trivial and the residuals are not uncorrelated (Thomaseth and Radde, 2016). This is often disregarded (see, e.g., (Degasperi et al., 2017)), which yields incorrect confidence intervals. In (ii), scaling parameters are introduced to replace the calibration curves. The scaling parameters are unknown and have to be inferred along with the dynamic parameters. While this increases the dimensionality of the optimization problem (see (Bachmann et al., 2011) for an example in which the number of parameters is doubled), the noise distribution is simple and the confidence intervals consistent. To address the dimensionality increase, Weber et al. (2011) proposed an approach for estimating the conditionally optimal scaling parameters given the dynamic parameters. This approach eliminated the scaling parameters, however, it is only applicable in the special case of additive Gaussian noise with known standard deviation. Unknown noise parameters and outlier-corrupted data (Maier et al., 2017) – as found in many applications – cannot be handled.

In this study, we propose a hierachical optimization approach which generalizes the idea of Weber et al. (2011). The proposed hierarchical approach allows for arbitrary noise distributions, with known and unknown noise parameters. For Gaussian and Laplace noise, we provide analytic solutions for the inner optimization problem, which boosts the computational efficiency. To illustrate the properties of the proposed approach, we present results for two models of JAK-STAT signaling and a model of RAF/MEK/ERK signaling.

## 2 Methods

In this section, we describe the considered class of parameter estimation problems and introduce a hierarchical optimization method for estimating the parameters of ODE models from relative data under different measurement noise assumptions.

### 2.1 Mechanistic modeling of biological systems

We considered ODE models of biological processes, 

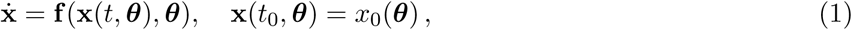

in which the time- and parameter-dependent state vector 
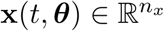
 represents the concentrations of the species involved in the process and the vector field 
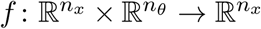
 determines how the concentrations evolve over time. The vector 
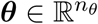
 denotes the parameters of the system, e.g., reaction rates. The initial conditions at time point *t*_0_ are given by the parameter-dependent function 
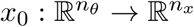
.

Experimental data provide information about observables 
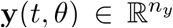
. These are obtained by the output function 
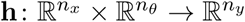
, which maps the states and parameters to the observables via

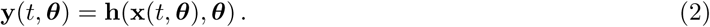

Due to experimental limitations the experimental data is noise corrupted, 

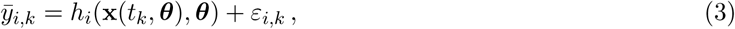

with *h_i_* denoting the *i*th component of the output function **h**, and indices *k* for the time point. In most applications, Gaussian noise is assumed, 
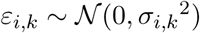
. For outlier-corrupted data, it was shown that the assumption of Laplace noise, *∈_i,k_*∼Laplace(0,*σ_i,k_*), yields more robust results (see (Maier et al., 2017) and references therein).

The measurements are collected in a dataset 
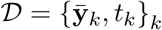
. The vector 
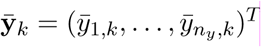
 comprises the measurements for the different observables. For the general case including different experiments and conditions, we refer to the Supplementary Information, Section 1.

### 2.2 Relative experimental data

Many experimental techniques provide data which are proportional to the measured concentrations. The scaling parameters are usually incorporated in **h**, defined in (2). Here, for simplicity and without loss of generality, we unplugged the scaling parameters from the function **h** and write

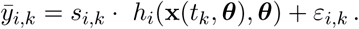

The scaling parameters *s_i,k_* and the noise parameters *σ_i,k_* are in the following combined in the matrices s and ***σ***, respectively. To distinguish the different parameter types, we refer to the parameters ***θ*** further as dynamic parameters. In the following, we present results for the case that the scaling *s_i_* and noise parameters *σ_i_* are the same for each time point, but differ between observables. The general case is presented in the Supplementary Information, Section 1.

### 2.3 Formulation of parameter estimation problem from relative data

We used maximum likelihood methods, a commonly used approach to calibrate mathematical model, to estimate the parameters from experimental data. The likelihood function is given by 

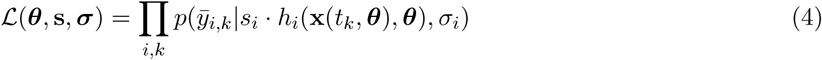

with *p* denoting the conditional probability of 
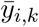
 given the observable *y_i,k_* = *s_i_* · *h_i_*(x(*t_k_*, ***θ***), ***θ***). This probability is for Gaussian noise

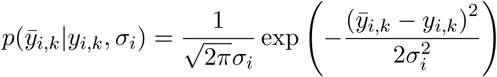

with standard deviation *σ_i_*, and for Laplace noise

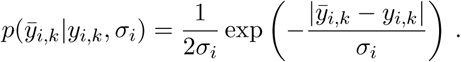

with scale parameter *σ_i_*.

#### 2.3.1 Standard approach to parameter estimation

For the standard approach, the dynamic parameters ***θ***, the scaling parameters s, and the noise parameters ***σ*** are estimated simultaneously. For numerical reasons, this is mostly done by minimizing the negative log-likelihood function,

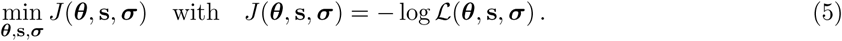

The parameters were combined as **q** = (**θ,s,σ**) and the optimization problem has the dimension: number of dynamic parameters *n_θ_* + number of scaling parameters *n_s_* + number of noise parameters *n_σ_*. We solved it using multi-start local optimization, a method which has previously been shown to be computationally efficient. In each iteration the objective function and its gradient were computed. If the objective function for this parameters fulfills certain criteria, e.g., the norm of the gradient was below a certain threshold, the optimization was stopped, otherwise the parameter was updated and the procedure was continued (Figure 1A).

**Figure 1:**
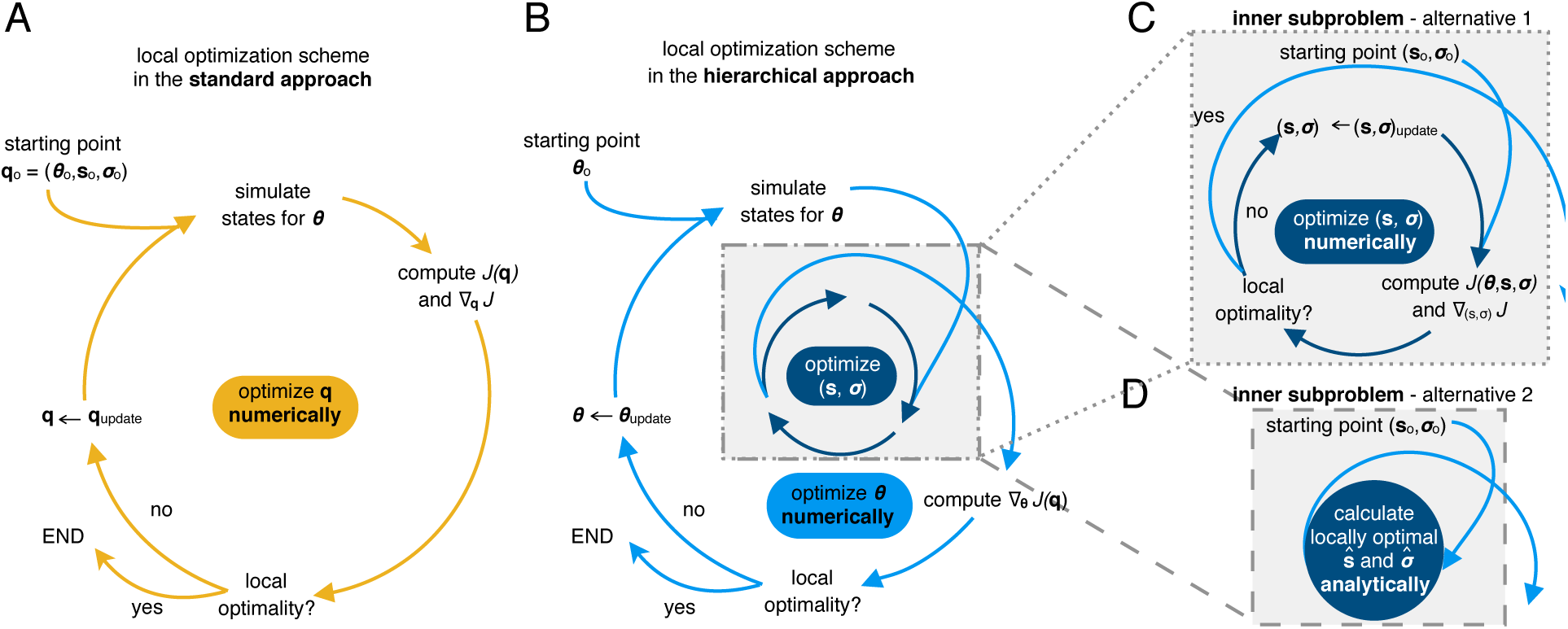
Visualization of standard and hierarchical optimization schemes. (**A**) Local optimization in the standard approach with parameters **q** = (***θ*,s,σ**). A single iteration includes the numerical simulation of the ODE model for ***θ***, the evaluation of the objective function and its gradient, the evaluation of local optimality and stopping criteria, and the termination of the local optimization or the updating of the parameters. (**B**) Outer local optimization in the hierarchical approach with parameters ***θ***. A single iteration includes the numerical simulation of the ODE model ***θ***, the evaluation of the objective function and its gradient with respect to ***θ*** using the results of the inner optimization problem, The iteration also includes the evaluation of local optimality and stopping criteria, and the termination of the local optimization or the updating of parameters. (**C,D**) Inner (local) optimization in the hierarchical approach to find the optimal scaling and noise parameter 
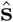
 and 
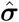
 for given dynamic parameters ***θ***. (**C**) Iterative local optimization to determine 
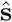
 and 
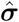
. This does not require the numerical simulation of the model. (D) Calculating optimal parameters 
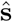
 and 
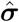
 using analytic expressions for common noise distributions.

#### 2.3.2 Hierarchical approach to parameter estimation

Since the optimization problem (5) often possess a large number of optimization variables and can be difficult to solve, we exploited its structure. Instead of solving simultaneously for **θ**,**s**, and **σ**, we considered the hierarchical optimization problem (Figure 1B)

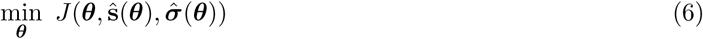

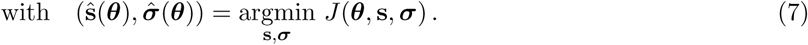

The inner problem (7) provides the optimal values 
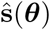
 and 
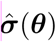
 of **s** and ***σ*** given ***θ***. These optimal values were used in the outer subproblem to determine the optimal value for ***θ*** denoted by 
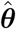
. It is apparent that a locally optimal point of the standard optimization problem (5) is also locally optimal for the hierarchical optimization problem (6,7), if the point is within the box constraints for the optimization.

The formulation (6) might appear more involved, however, it possesses several properties which might be advantageous:

i. The individual dimensions of the inner and outer subproblems (6,7) are lower than the dimension of the original problem (5).
ii. The optimization of the inner subproblem does not require the repeated numerical simulation of the ODE model.
iii. For several noise models, e.g., Gaussian and Laplace noise, the inner subproblem can be solved analytically.

If (iii) holds, the scaling parameters s and also the noise parameters ***σ*** can be calculated directly and the amount of parameters that need to be optimized iteratively reduces to *n_θ_* (Figure 1C,D). In the following two sections, the analytic expressions for the Gaussian and Laplace noise are derived. For this, let observable index *i* be arbitrary but fixed.

## Analytic expressions for the optimal scaling and noise parameters for Gaussian noise

In this study, we evaluated the scaling and noise parameters for Gaussian noise analytically. To derive the analytic expression for the optimal parameters, we exploited that the objective function for Gaussian noise,

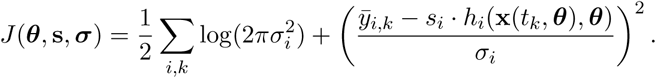

is continuously differentiable, and that the gradient of *J* at a local minimum is zero. For the inner subproblem this implies

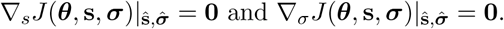

These equations can be solved analytically (see Supplementary Information, Section 1), which yields

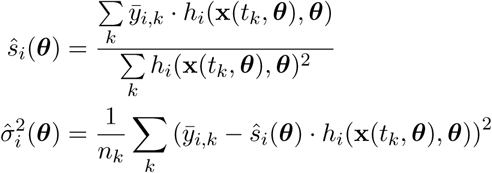

with number of time points *n_k_*. Consistent with the structure of the hierarchical problem (6), both formulas depend only on the dynamic parameters ***θ***.

In many studies (e.g., (Bachmann et al., 2011)), observation functions of the form 
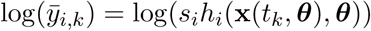
 are used. In the Supplementary Information, Section 2, we provide a derivation of the corresponding optimal parameters.

## Analytic expressions for the optimal scaling and noise parameters for Laplace noise

For Laplace noise the negative log-likelihood function is

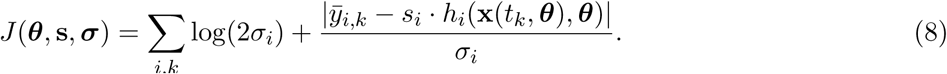

This objective function is continuous but not continuously differentiable. In this case, a sufficient condition for a local minimum is that the right limit value of the derivative is negative and the left limit value is positive. The derivative of (8) with respect to *s_i_* can be written as

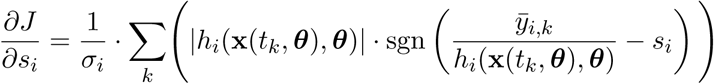

As *σ_i_* is positive, the locations of kinks in the objective function and the corresponding jumps in the derivative are independent of *σ_i_* (Figure 2). Accordingly, the problem of finding 
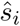
 reduced to checking the signs of the derivative before and after the jump points 
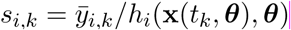
. We sorted *s_i,k_* in increasing order and evaluated the derivatives at the midpoints between adjacent jumps, a procedure which is highly efficient as the ODE model does not have to be simulated. Given 
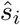
, the noise parameter 
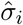
 follows from the work of Norton (1984)

**Figure 2:**
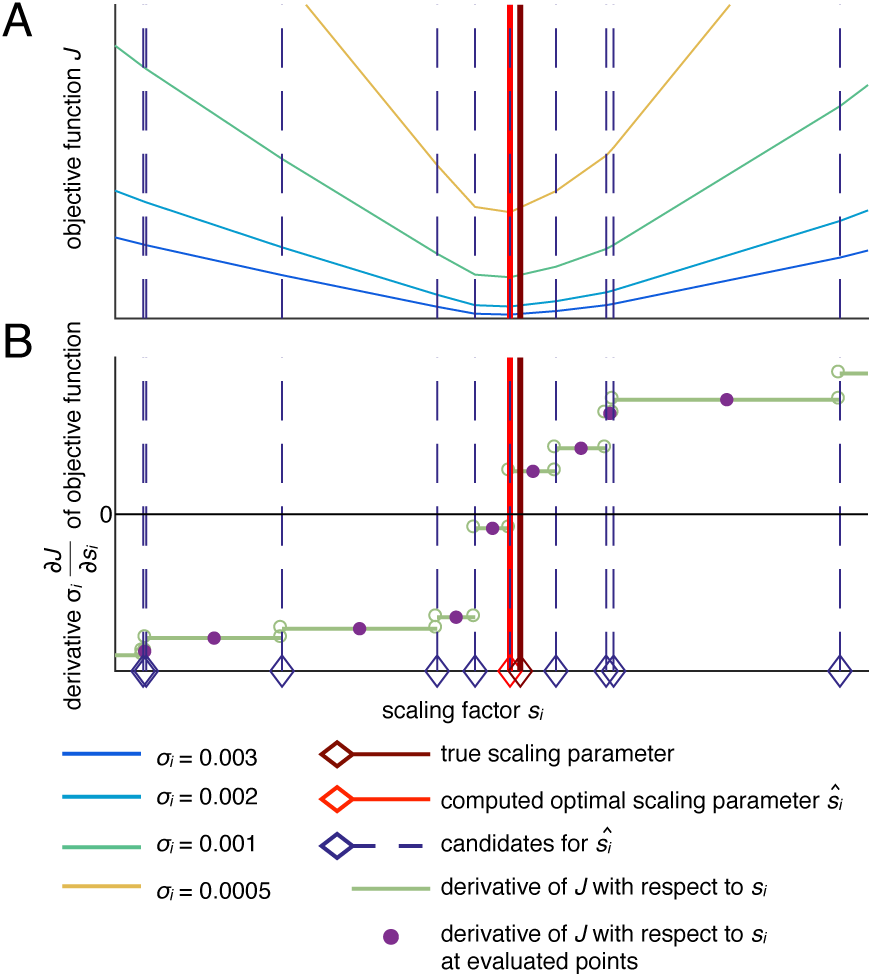
Illustration of the computation of an optimal scaling parameter 
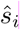
 for Laplace noise. (**A**) Objective function *J* for different values of *σ_i_*, showing that the kinks indicated by the dashed lines are independent of that value. (**B**) Derivative of the objective function with respect to the scaling parameter which is not defined at the kinks. The light red and dark red lines indicate the computed scaling parameter and the true optimal scaling parameter, respectively.

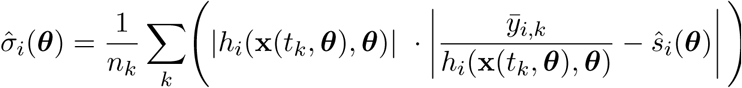

Both derived formulas depend only on the dynamic parameters ***θ***, in consistence with the structure of the hierarchical problem (6). In summary, we reformulated the original optimization problem (5) as a hierarchical optimization problem (6,7), and provided an analytic solution to the inner subproblem (7) for several relevant cases. Using the analytic solutions, the kinetic parameters can be inferred by solving a lower-dimensional problem.

## 3 Results

To study and compare the performance of parameter estimation from relative data using the standard approach and our hierarchical approach, we applied both to three published estimation problems.

### 3.1 Models and experimental data

The considered models describe biological signaling pathways, namely, the JAK-STAT (Swameye et al., 2003; Bachmann et al., 2011) and the RAF/MEK/ERK signaling pathway (Fiedler et al., 2016).

#### 3.1.1 JAK-STAT signaling I

The first application example we considered is the model of Epo-induced JAK-STAT signaling introduced by Swameye et al. (2003) (Figure 3A). Epo yields the phosphorylation of signal transducer and activator of transcription 5 (STAT5), which dimerizes, enters the nucleus to trigger the transcription of target genes, gets dephosphorylated, and is transported to the cytoplasm. We implemented the model which describes the phosphorylated Epo receptor concentration as a time-dependent spline (Schelker et al., 2012). For further details on the model, we refer to Supplementary Information, Section 4.1.

**Figure 3:**
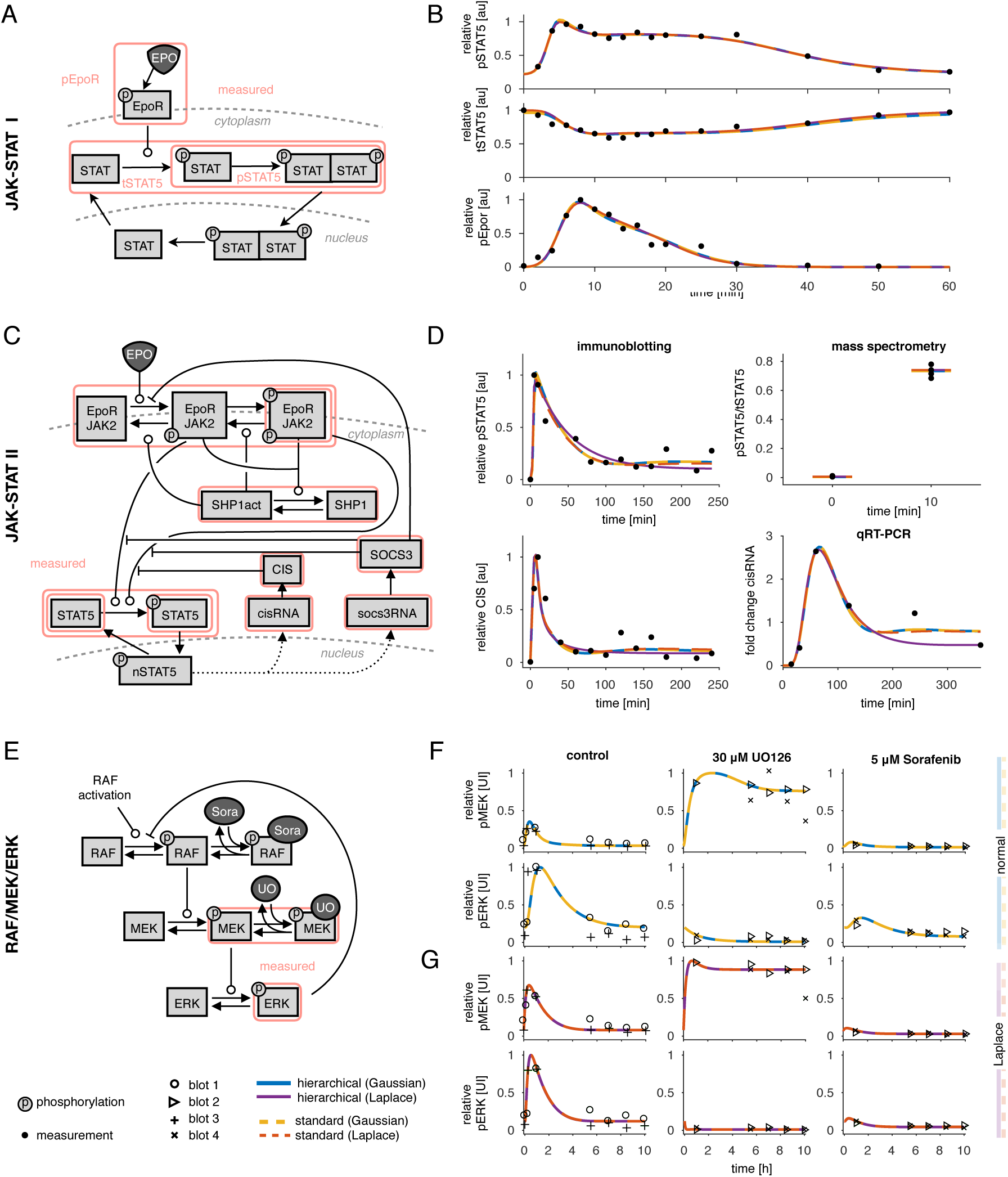
Models and experimental data. (**A,B**) JAK-STAT I. (**A**) Illustration of the model according to Swameye et al. (2003). Arrows represent biochemical reactions, and the observables of the model used are highlighted by boxes. (**B**) Experimental data and fitted trajectories for the best parameter found with multi-start local optimization with 100 starts. The results are shown for the standard (dotted lines) and hierarchical (solid lines) approach for optimization for Gaussian and Laplace noise. (**C,D**) JAK-STAT II. (**C**) Illustration of the model according to Bachmann et al. (2011). (**D**) Experimental data and fitted trajectories for the best parameter found with multi-start local optimization for 200 starts. 33 out of 541 data points are shown. (**E-G**) RAF/MEK/ERK. (**E**) Illustration of the model according to Fiedler et al. (2016). (**F,G**) Experimental data and fitted trajectories for the best parameter found with multi-start local optimization for 500 starts. different markers indicate the different blots. The data is scaled according to the estimated scaling parameters, yielding different visualizations for different parameters, as obtained with the Gaussian and the Laplace noise assumption. (**F**) Fitted trajectories for Gaussian noise for the standard (dotted line) and hierarchical (solid line) approach for optimization. (**G**) Fitted trajectories for Laplace noise.

The model parameters were estimated using immunoblotting data for the phosphorylated Epo receptor (pEpoR), phosphorylated STAT5 (pSTAT5), and the total amount of STAT5 in the cytoplasm (tSTAT5) (Figure 3B). Experimental data are available for 16 different time points. Since immunoblotting only provides relative data, the scaling parameters for the observables need to be estimated from the data. As proposed by Schelker et al. (2012), the scaling parameter for pEpoR has been fixed to avoid structural non-identifiabilities (Raue et al., 2009). This yields *n_θ_* = 11 dynamic parameters (see Supplementary Information, Section 4.1), *n_s_* = 2 scaling parameters, and *n_σ_* = 3 noise parameters.

#### 3.1.2 JAK-STAT signaling II

The second application example is the model of JAK-STAT signaling introduced by Bachmann et al. (2011). This model provides more details compared to the previous one. It includes, for instance, gene expression of cytokine-inducible SH2-containing protein (CIS) and suppressor of cytokine signaling 3 (SOCS3), and possesses more state variables and parameters (Figure 3C).

The model parameters were estimated using immunoblotting, qRT-PCR, and quantitative mass spectrometry data (Figure 3D and Supplementary Information, Figure S4). To model the observables Bachmann et al. (2011) used *n_s_* = 43 scaling parameters, and *n_σ_* = 11 noise parameters, yielding *n_θ_* = 58 remaining parameters. Some scaling and noise parameters are shared between experiments and some are shared between observables. For this model, most of the observables were compared at the log_10_ scale (see Supplementary Information, Section 4.2).

#### 3.1.3 RAF/MEK/ERK signaling

The third application example we considered is the model of RAF/MEK/ERK signaling introduced by Fiedler et al. (2016). The model describes the phosphorylation cascade and a negative feedback of phosphorylated ERK on RAF phosphorylation (Figure 3E).

Fiedler et al. (2016) collected Western blot data for HeLa cells for two observables, phosphorylated MEK, and phosphorylated ERK, with four replicates at seven time points (Figure 3F,G). Each observable and replicate was assumed to have different scaling and noise parameters, yielding 16 additional parameters (Figure 4A).

**Figure 4:**
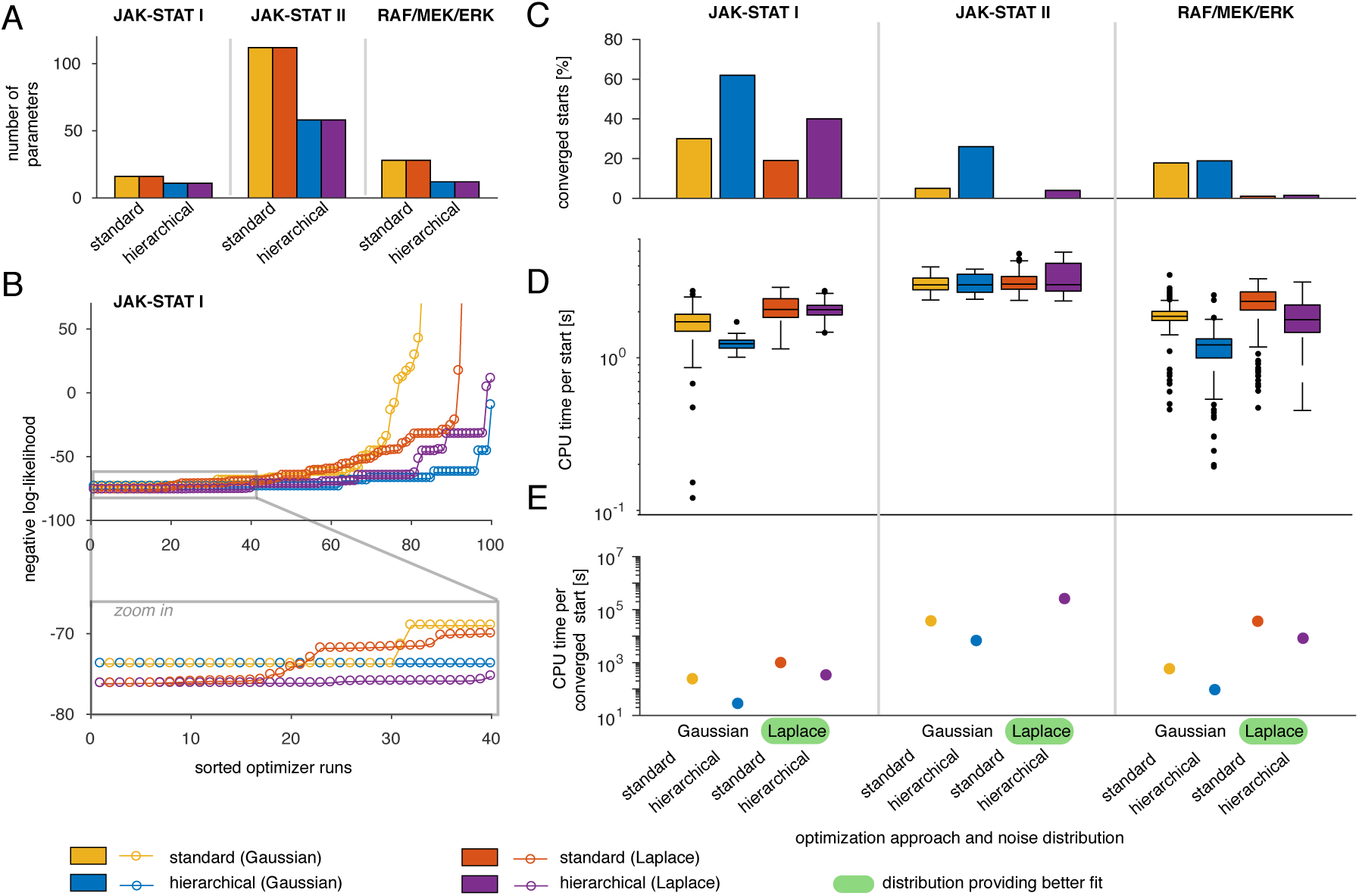
Evaluation of the standard and hierarchical approach for three application examples. (**A**) Number of parameters which need to be optimized numerically. (**B**) Likelihood waterfall plot for the JAK-STAT model I. The ascendingly sorted negative log-likelihood values are shown for both approaches (standard and hierarchical) and noise distributions (Gaussian and Laplace). (**C-E**) Comparison of the two optimization approaches and two noise distribution for the three models. The noise model with the better likelihood function is highlighted in the label. (**C**) Percentage of converged starts over all performed local optimizations. (**D**) Boxplot for the CPU time needed per start. (**E**) CPU time needed per converged start.

### 3.2 Evaluation of the approaches

We performed parameter estimation for the application examples using the standard and the hierarchical approach. For each example, the case of Gaussian and Laplace noise was considered. The resulting optimization problems were solved with the MATLAB toolbox PESTO (Stapor et al., 2017), using multi-start local optimization, an approach which was previously found to be computationally efficient and reliable (Raue et al., 2013). Initial points were sampled uniformly within their parameter boundaries and local optimization was performed using the interior point method implemented in the MATLAB function `fmincon.m.` Numerical simulation and forward sensitivity analysis for gradient evaluation was performed using the MATLAB toolbox AMICI (Fröhlich et al., 2017), which provides an interface to CVODES (Serban and Hindmarsh, 2005). To improve convergence and computational efficiency, log_10_-transformed parameters were used for the optimization.

#### 3.2.1 Qualitative comparison of optimization approaches for different noise distributions

As the standard and hierarchical approach should in principle be able to achieve the same fit, we first studied the agreement of trajectories for the optimal parameters. We found that they coincide for the JAK-STAT model I and the RAF/MEK/ERK model, indicating that the hierarchical approach is able to find the same optimal value as the standard approach (Figure 3B,F,G). Also the best likelihood values which were found for these two models by the two approaches coincide (Figure 4B and Supplementary Information, Figure S5). Only for the JAK-STAT model II for the case of Laplace noise, the fitted trajectories deviate (Figure 3D). Insertion of the optimum found by the hierarchical approach in the objective function of the standard approach revealed that the standard approach missed the optimal point (SupplementaryInformation, Figure S3). As expected, there are di erences between the results obtained with Gaussian and Laplace noise, which is visible in the trajectories and the corresponding likelihood values. Interestingly, for each model the likelihood values achieved using Laplace noise were better than for Gaussian noise (Supplementary Information, Figure S1C). This indicates that the Laplace distribution with its heavier tail is more appropriate than the Gaussian distribution for the considered estimation problems.

#### 3.2.2 Convergence of optimizers

As the performance of multi-start local methods depends directly on the convergence of the local optimizers, we assessed for how many starting points the local optimizer reached the best objective function value found across all runs. This was done by studying the likelihood waterfall plots (Figure 4B). We found that the proposed hierarchical approach achieved consistently a higher fraction of converged starts than the standard approach (Figure 4C). Local optimization using the hierarchical approach converged on average in 25.38% of the runs while the standard approach converged on average in 12.13% of the runs.

The application examples vary with respect to the total number of parameters and in the number of parameters which correspond to scaling or noise parameters (Figure 4A). While for the JAK-STAT model I only five parameters could be optimized analytically, for the JAK-STAT model II almost half of the parameters correspond to scaling or noise parameters. Interestingly, even when the dimension of the optimization problem was only reduced by few parameters, we observed a substantial improvement of the convergence (Figure 4C).

#### 3.2.3 Computational efficiency

As computation resources are often limiting, we finally analyzed the computation time per converged start. We found that on average, the computation time per start was lower for the hierarchical approach than for the standard approach (Figure 4D). In combination with the improved convergence rate, this resulted in a substantially reduced computation time per converged start, aka a start which reach the minimal value observed across all starts (Figure 4E). Given a fixed computational budget, the hierarchical approach achieved on average 5.52 times more optimization runs which reached the best objective function values than the standard approach.

In summary, the application of our hierarchical approach to parameter estimation from relative data to the models shows consistently that our approach yields parameter values of the same quality as the standard method, while achieving better convergence and reducing the computation time substantially.

## 4 Conclusion

The statistically rigorous estimation of model parameters from relative data requires non-standard statistical models (Degasperi et al., 2017) or scaling parameters (Raue et al., 2013). Unfortunately, the former is not supported by established toolboxes and the latter increases the dimensionality of the estimation problem. In this manuscript, we introduced a hierarchical approach which avoids the increase of dimensionality and is applicable to a broad range of noise distributions. For Gaussian and Laplace noise we provided analytic expressions. The approach can be used for combinations of relative and absolute data, and for different optimization methods, including least-squares methods or global optimization methods such as particle swarm optimization (Vaz and Vicente, 2009) (see Supplementary Information, Figure S2).

We evaluated the performance of our hierarchical approach and compared it to the standard approach for three models, which vary in their complexity. For all applications, we found that our hierarchical approach yielded fits of the same or better quality. In addition, convergence was improved and the computation time was shortened substantially. We demonstrated that our approach can also be used when relative and absolute data are modeled together in an experiment, and when several observables or experiments share scaling and/or noise parameters. This renders our approach applicable to a wide range of mathematical models studied in systems and computational biology. We provided a generic implementation of the objective function for the hierarchical approach for Gaussian and Laplace noise. The objective function is provided in the Supplementary Information (along with the rest of the code) and included in the MATLAB toolbox PESTO (Stapor et al., 2017).

In addition to the scaling and noise parameters, also other parameters which only contribute to the mapping from the states to the observables, could be optimized analytically. This includes o set parameters, which are used to model background intensities or unspecific binding. Extending our approach to also calculate these parameters analytically would decrease the parameters in the outer optimization even more.

We employed forward sensitivities for the calculation of the objective function gradient. However, it has been shown that for large-scale models with a high number of parameters, adjoint sensitivities can reduce the computation time needed for simulation (Fröhlich et al., 2017). Thus, a further promising approach would be the combination of both complementary approaches for the handling of large-scale models.

To summarize, employing our hierarchical approach for optimization yielded more robust results and speed up the computation time. This renders the approach valuable for estimating parameters from relative data. The proposed approach might facilitate the handling of large-scale models, which possess many measurement parameters.

## Funding

This work has been supported by the European Union’s Horizon 2020 research and innovation program under grant agreement no. 686282

## 1 General formula for analytic scaling and noise parameters

In the main manuscript, we covered experimental data sets which have different time points. Here, we provide the derivation of the expressions for the general case, in which the experimental data also comprise different replicates, experiments, and conditions, e.g., varying drug doses. We considered that the ODE system also depends on an input 
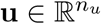
,

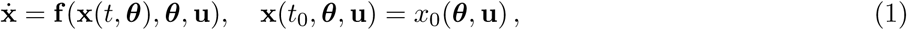

thus, 
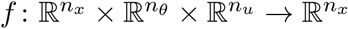
 which also affects the mapping to the observables

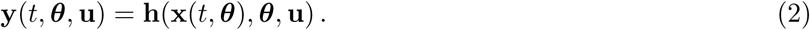

The experimental data is then given by

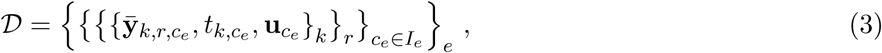

including all indices for time point *k*, replicate *r*, experiment-specific condition *c*_*e*_, and experiment *e*. The indices *I*_*e*_ indicate which conditions correspond to a certain experiment. The measurements are mapped to the states by

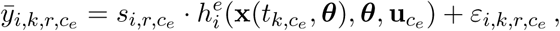

with 
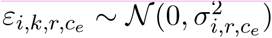
 or 
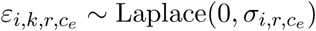
 and 
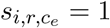
 for absolute measurements. Also, the structure of the mapping from states to observables might be experiment-specific. The negative log-likelihood is given by

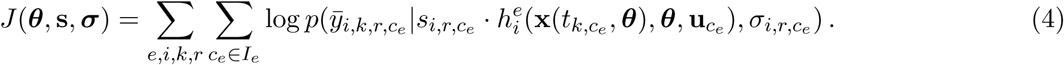

In the main manuscript, we presented the analytic formulas for the case that each observable and corresponding replicate has different scaling and noise parameters, but that these parameters do not change between conditions and time points. A more general formula is provided in the following, covering, e.g., the case that replicates share the same scaling parameters, but observables do not. This can be easily generalized to also include variability between time points.

### 1.1 Gaussian noise

The general objective function under Gaussian noise is given by

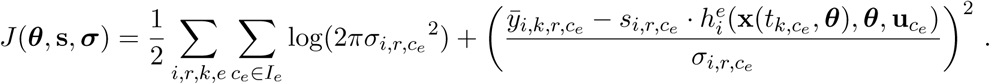

To define which replicates, observables, and experiments share a scaling or noise parameter, we define

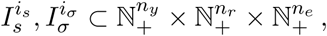

for *i*_*s*_ = 1, …, *n*_*s*_ and *i*_σ_ = 1, …, *n_σ_*. The number of replicates is denoted by n_r_ and the number of experiments by *n*_*e*_. This means, all scaling parameters 
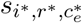
 for which the indices (i^*^, r^*^, e^*^) are part of the same group *I*_s_ share the same scaling parameters. This yields *n*_*s*_ different scaling parameters that are estimated from the data. For this we denote 
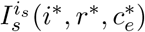
 the group which includes the indices 
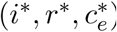
. This is analogously for the noise parameters. The derivative of the objective function with respect to a scaling parameter thus reads

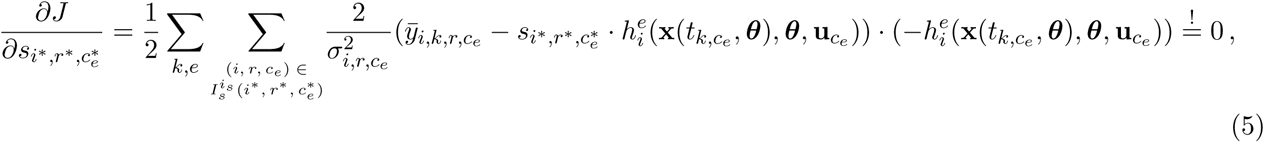

and was set to zero to obtain the analytic expression for the optimal scaling parameter. The solution does not depend on the noise parameters if 
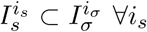
 and we solve the equation with respect to 
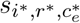
 to obtain the optimal value

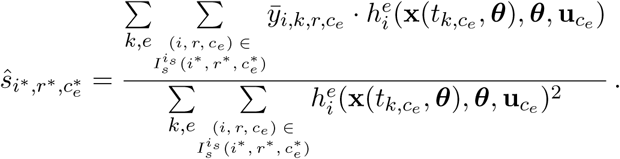

For the noise parameters, we need

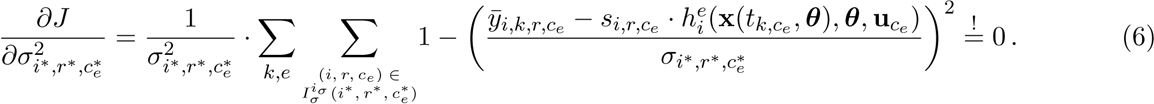

We write

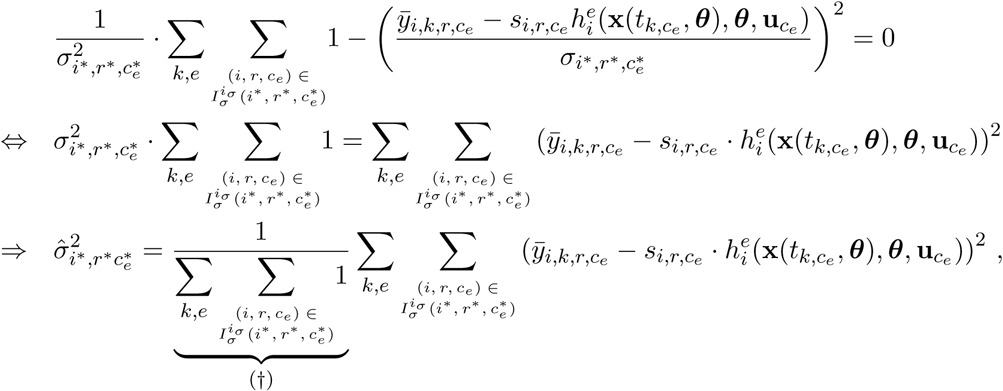

in which (†), the nominator, is simply the number of observations in which 
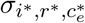
 appears. In some cases, for instance if all experiments share the same scaling parameter, we neglected the superscript *e*.

The gradient used for optimization is given by

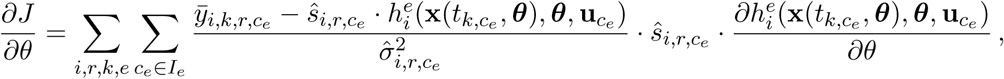

for which 
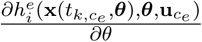
 is obtained by forward sensitivity equations employed in AMICI, and,

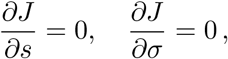

which holds due to (5) and (6). The Hessian with respect to the dynamic parameters is

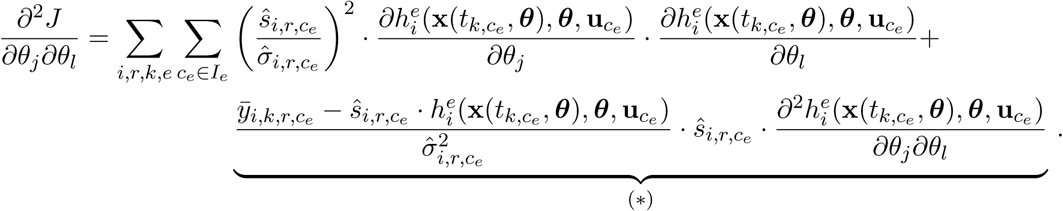

For the remaining parameter, the Hessian is zero. We implemented an approximation of the Hessian neglecting the terms (*) that include higher-order sensitivities.

### 1.2 Laplace noise

For Laplace noise, the expression for the optimal scaling and noise parameters can be generalized analogously. The objective function for the general case is

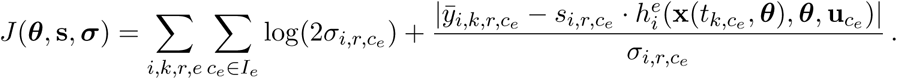

The derivative with respect to a scaling parameter is

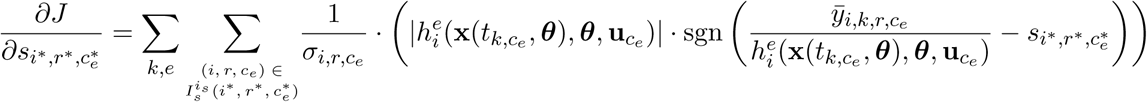

with jump points

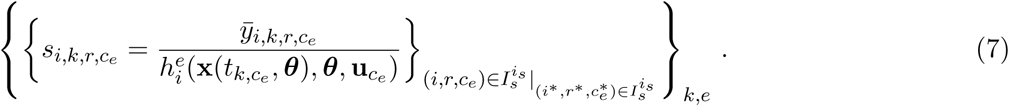

These jump points are the candidates for the optimal scaling parameter and the candidate for which the sign of the derivative changes is chosen. For the optimal noise parameter we have

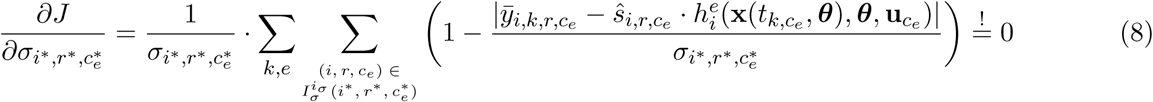

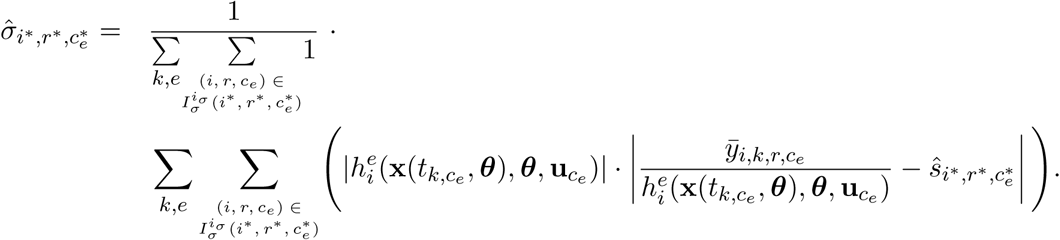

The gradient used for optimization is given by

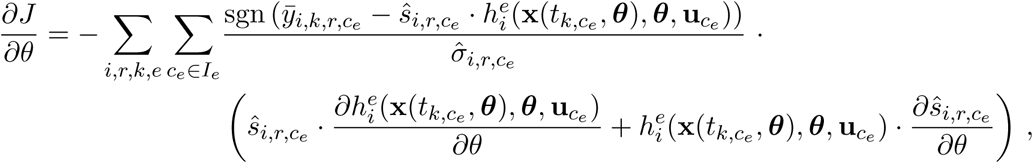

for which 
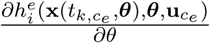
is obtained by forward sensitivity equations employed in AMICI, and 
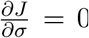
, which holds due to (8).

## 2 Comparison of data and simulation at a logarithmic scale

In the main manuscript and Supplementary Information, Section 1, we provided the formulas for the comparison of data and simulation on a linear scale. However, sometimes it might be more appropriate to compare experimental data and simulation on a logarithmic scale.

### 2.1 Gaussian noise

For Gaussian noise, the objective function for the comparison on the logarithmic scale is given by

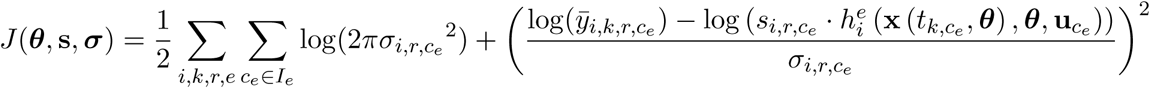

Thus, the derivative with respect to the scaling parameters is

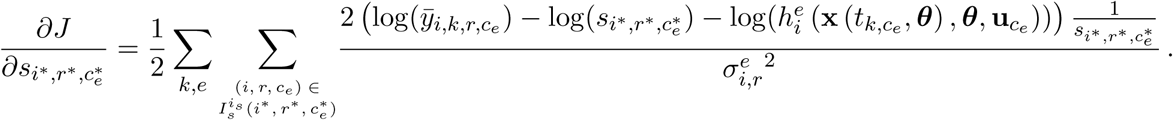

This yields the formula for the optimal scaling parameters

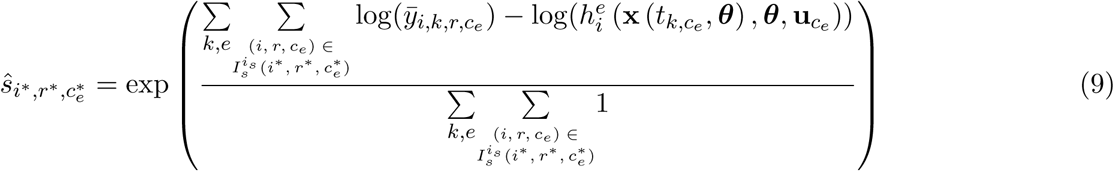

and

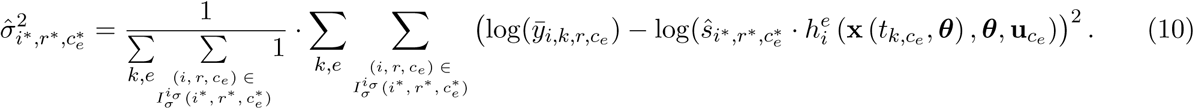

The gradient used for optimization is given by

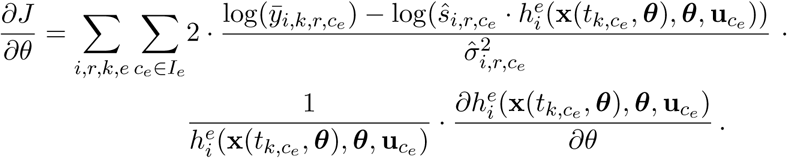

If the data is compared at log_10_ scale, as, e.g., for the JAK-STAT signaling model proposed by Bachmann et al. (2011), the negative log-likelihood function reads

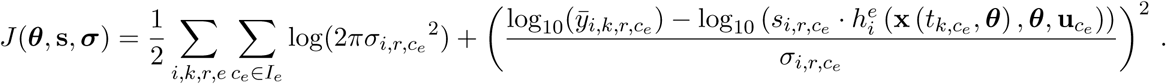

The optimal scaling parameters here are the same as when using the natural logarithm (9). For the optimal noise parameters the log is replaced by log_10_ in (10).

### 2.2 Laplace noise

For the Laplace distribution including the logarithmic comparison

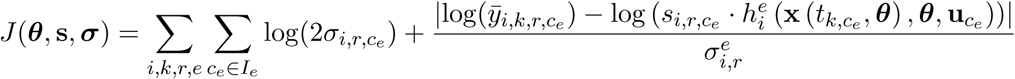

the same procedure can be applied for the logarithmic scale as for the linear scale, with the same set of candidate scaling parameters (7) as for the linear scale. However, one has to pay attention to adapt the derivative properly, for which the change of signs is checked. The optimal noise parameters then is given by

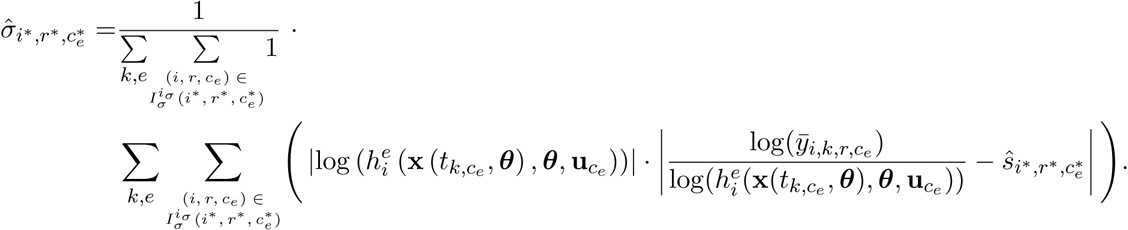

The gradient used for optimization is given by

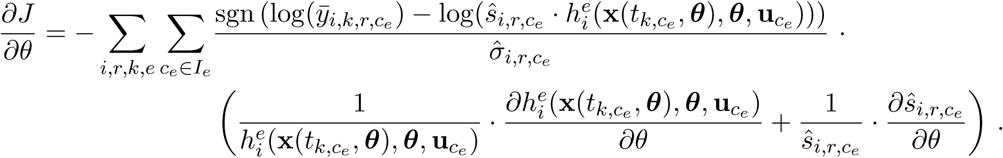

## 3 Implementation

We implemented the log-likelihood function and the analytic calculation of the scaling and noise parameters in easy-to-use MATLAB functions. The log-likelihood function is provided in `loglikelihoodHierarchical.m`, which provides the log-likelihood value, the gradient of the log-likelihood function with respect to the dynamic parameters, and in the case of Gaussian noise also an approximation to the Hessian by neglecting second-order derivatives. The functions and examples are incorporated in the toolbox PESTO (Stapor et al., 2017)and can be found on GitHub: http://github.com/ICB-DCM/PESTO. The simulated observables, their sensitivities, the experimental data, and the specification of measurement noise, scale of comparison between simulation and data, and shared parameters needs to be supplied by the user.

For our analysis, we employed the toolbox AMICI (Fröhlich et al., 2017) for the simulation of the system and the simulation of the sensitivities, and the toolbox PESTO (Stapor et al., 2017) for the estimation of the parameters.

## 4 Models and experimental data

In the following, we provide the details of the mathematical models. The considered models vary in their number of parameters (Figure 6A), number of data points that are used to calibrate the models (Figure S1A), and number of states of the underlying ODE system (Figure S1B).

### 4.1 JAK-STAT signaling I

For the first model, we used the model introduced by Schelker et al. (2012), which is defined by the ODE system

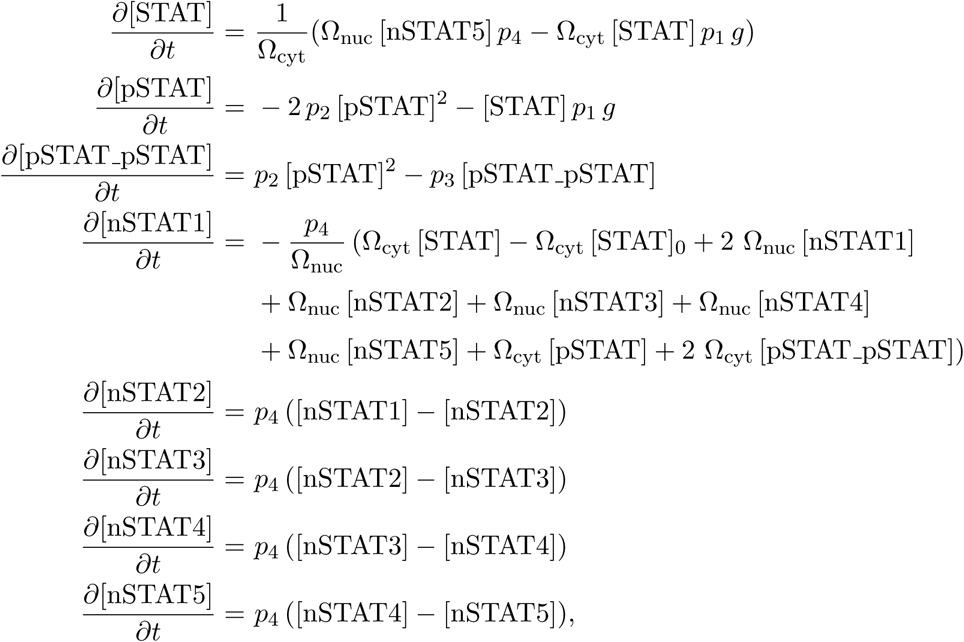

**Supplementary Figure S1:**
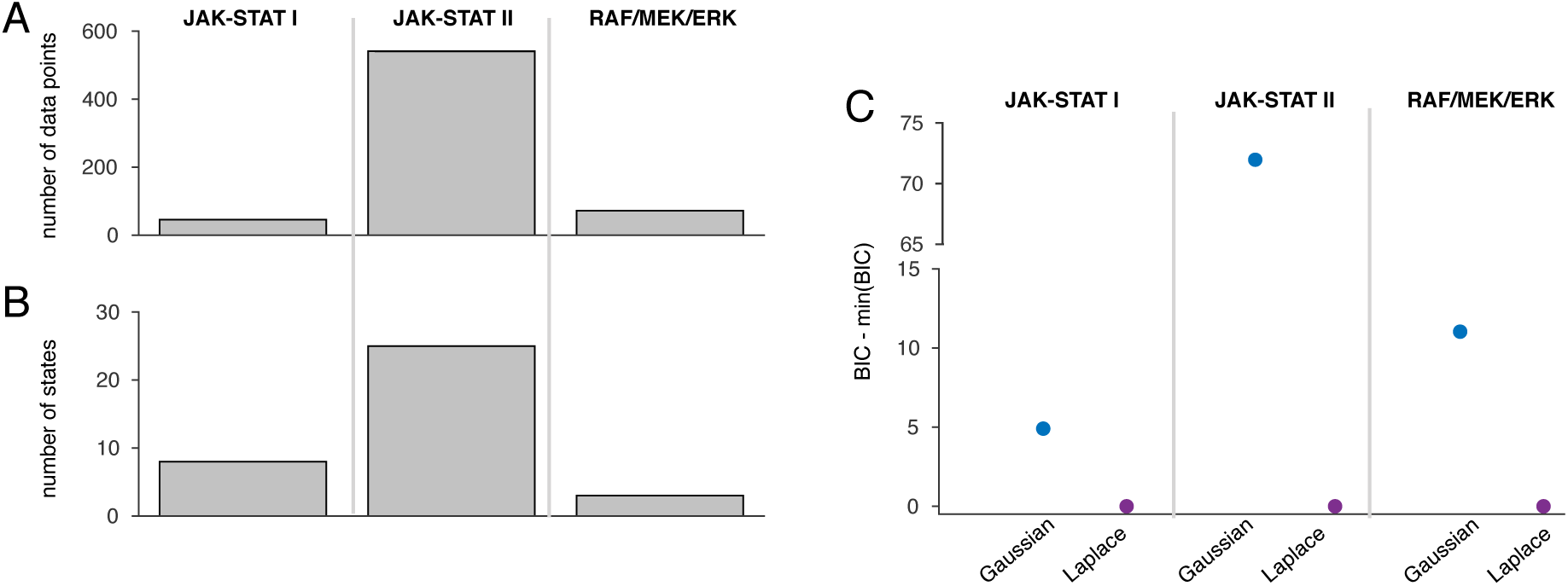
Comparison of the three models. (A) Number of experimental data points used to calibrate the models. (B) Number of states *n*_*x*_. (C) Comparison of Gaussian and Laplace noise for the three models based on the Bayesian Information Criterion (BIC) (Raftery, 1999), which rewards high likelihood values and penalizes high number of parameters.

with kinetic parameters *p*_1_,…, *p*_4_. The brackets indicate the concentrations of the corresponding species. The initial conditions are given by

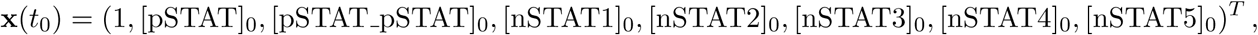

for which the initial condition for STAT is set to 1 in order to remove structural non-identifiabilities (Schelker et al., 2012). The states nSTAT1, …, nSTAT5 are intermediate steps, resulting from a linear chain approximation to model the delay of STAT binding to the DNA in the nucleus. The volumes of the cytoplasm and nucleus are denoted by Ω_cyt_ = 1.4 pl and Ω_nuc_ = 0.45 pl, respectively (Raue et al., 2009).

The observables are defined by *y*_1_ for total concentration of phosphorylated STAT in the cytoplasm (pSTAT), *y*_2_ for the total concentration of STAT in the cytoplasm (tSTAT), and *y*_3_ for the phosphorylated Epo receptors (pEpoR) (see Figure 3A in the main manuscript). They are linked to the states of the system via

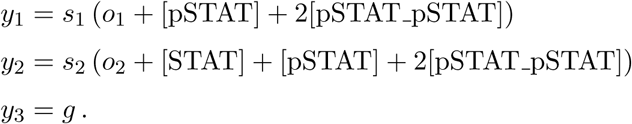

The concentration of Epo receptors is modeled as time-dependent cubic spline function *g* with parameters *sp*_1_, …, *sp*_5_, which are also estimated from the data. The parameters *o*_1_ and *o*_2_ define the to offsets needed to model the background noise. The model comprises the parameters q = (*p*_1_, *p*_2_, *p*_3_, *p*_4_, sp_1_, *sp*_2_, *sp*_3_, *sp*_4_, *sp*_5_, *o*_1_, *o*_2_, *s*_1_, *s*_2_, *σ*_1_, *σ*_2_, *σ*_3_)^T^, for which *θ*= (*p*_1_, *p*_2_, *p*_3_, *p*_4_, *sp*_1_, *sp*_2_, *sp*_3_, *sp*_4_, *sp*_5_, *o*_1_, *o*_2_) was optimized in the outer optimization problem of the hierarchical approach. The scaling parameters s = (*s*_1_, *s*_2_) and noise parameters= (σ_1_, σ_2_, σ_3_) for observables *y*_1_, *y*_2_, and *y*_3_, respectively, were optimized in the inner optimization problem. The subscript for these parameters indicates the observable. We neglected indices *r*, *e*, and *c*_*e*_, since only one experiment, replicate, and condition is considered. The parameter boundaries for the optimization are given

**Supplementary Figure S2:**
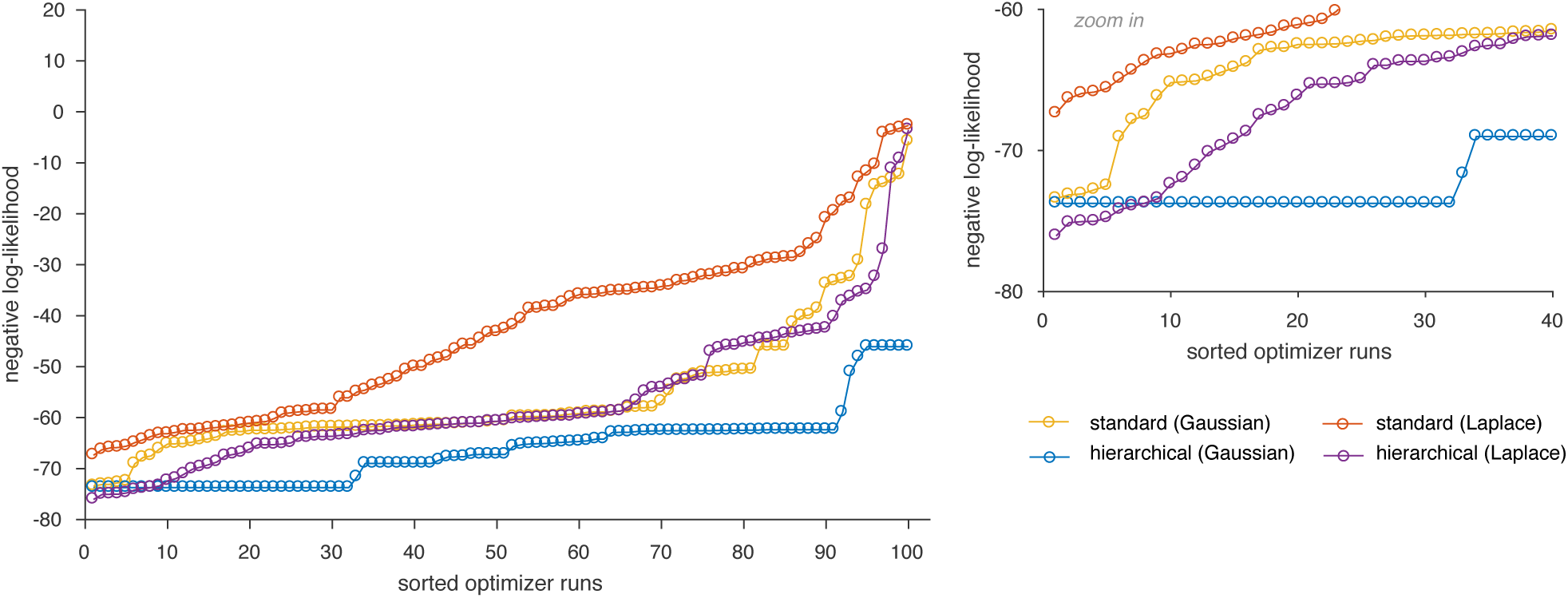
Likelihood waterfall plot for JAK-STAT signaling I using particle swarm optimization.

by

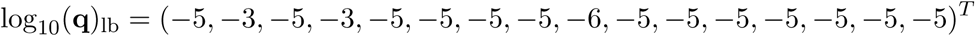

for the lower bound and

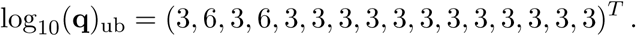

for the upper bound (Maier et al., 2017). We performed 100 optimizations, starting from randomly drawn parameter values. The starting points for the dynamic parameters were the same for both optimization approaches.

The comparison between the two noise assumptions revealed that the Laplace noise is more appropriate. However, the difference in BIC values was below 10, which indicates that the improvement was not substantial (Figure S1C) (Kass and Raftery, 1995).

To evaluate the possibility of using the hierarchical optimization also within global optimization, we repeated the analysis using an particle swarm algorithm (Vaz and Vicente, 2009). This method does not need gradient information and has been shown to outperform other global optimization methods (Vaz and Vicente, 2009). The waterfall plots are shown in Figure S2. Interestingly, only the hierarchical optimization for the Gaussian noise was able to find the same optimum as the deterministic optimization. For the other settings the convergence suffered. However, as for the optimization with `fmincon`, the hierarchical approach was superior to the standard approach and the Laplace noise fitted the data better than the Gaussian noise.

### 4.2 JAK-STAT signaling II

The ODE system for JAK-STAT signaling model II is given by (Bachmann et al., 2011)

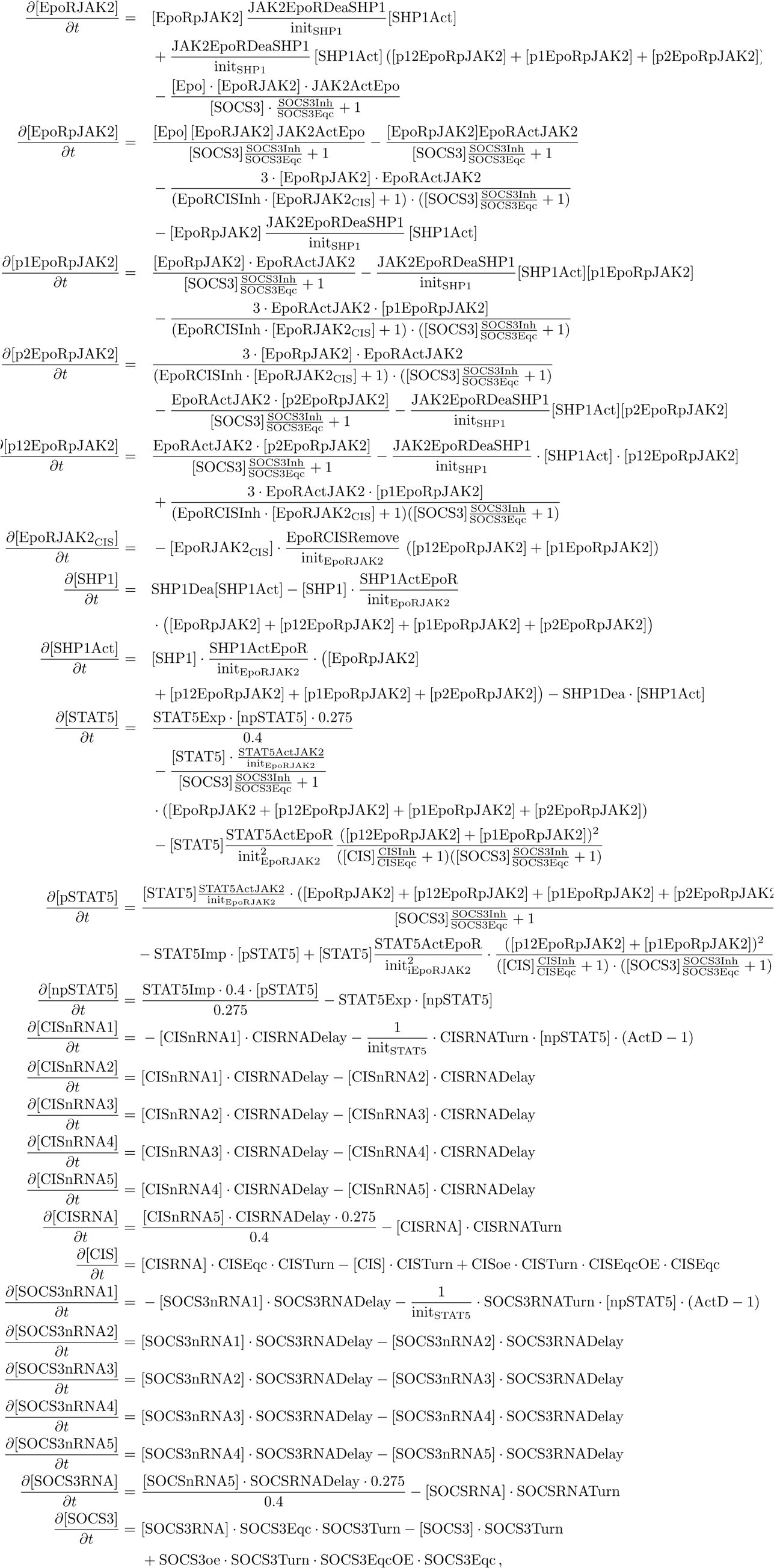

with condition-specific initial conditions (see Table S1) denoted by 
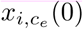
 for observable index *i* under condition indexed by *c*_*e*_:

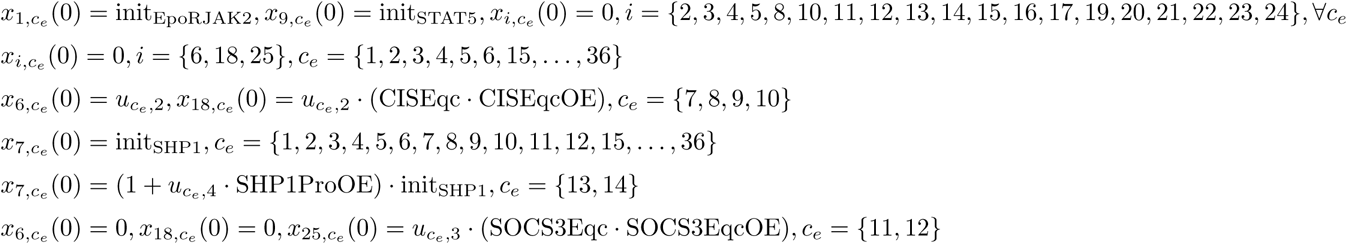

The observables are given by

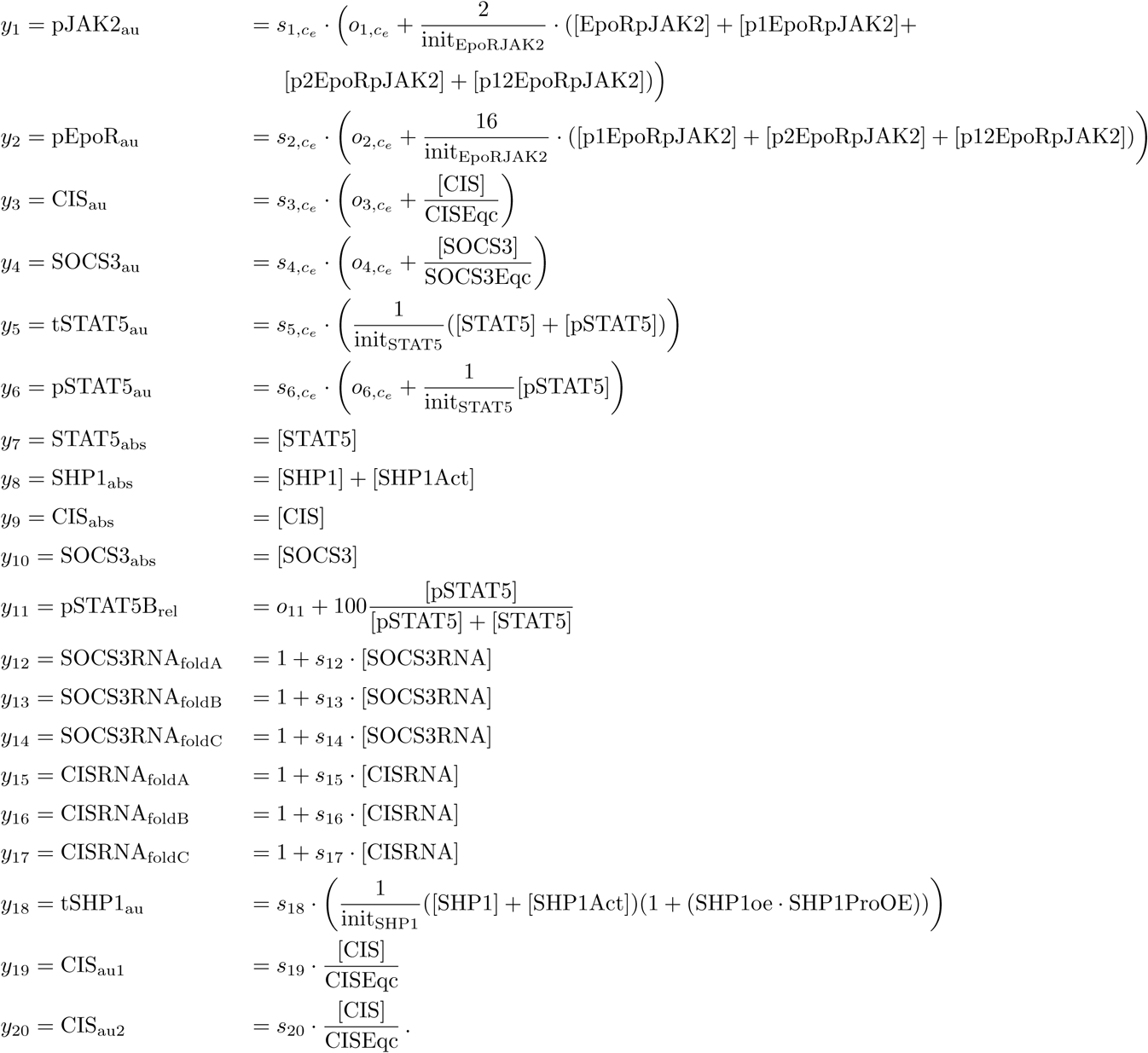

The parameters *θ* are

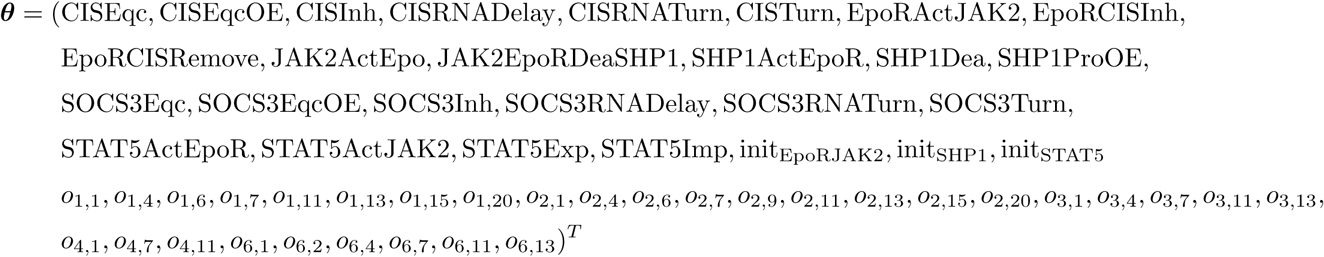

with *n*_θ_ = 58. For experiment SHP1oe (*e* = 9), the parameter init_SHP1_ was replaced by init_SHP1_ · (1 + (SHP1oe · SHP1ProOE)) in the model equations. For the notation of the offset, scaling, and noise parameters, we neglected the index *r*, since these parameters are shared for the replicates. The first subscript indicates the observable, and the second the condition. However, all conditions belonging to the same experiment share the scaling and offset parameters and thus the parameters are only listed for the first condition of each experiment. The experiments and corresponding condition indices are summarizes in Table S1. For simplicity, we note the scaling parameters as vector s which contains only the unique parameters *s_i_*,*c_e_* which need to be estimated from the data. Thus, it is

**Supplementary Figure S3:**
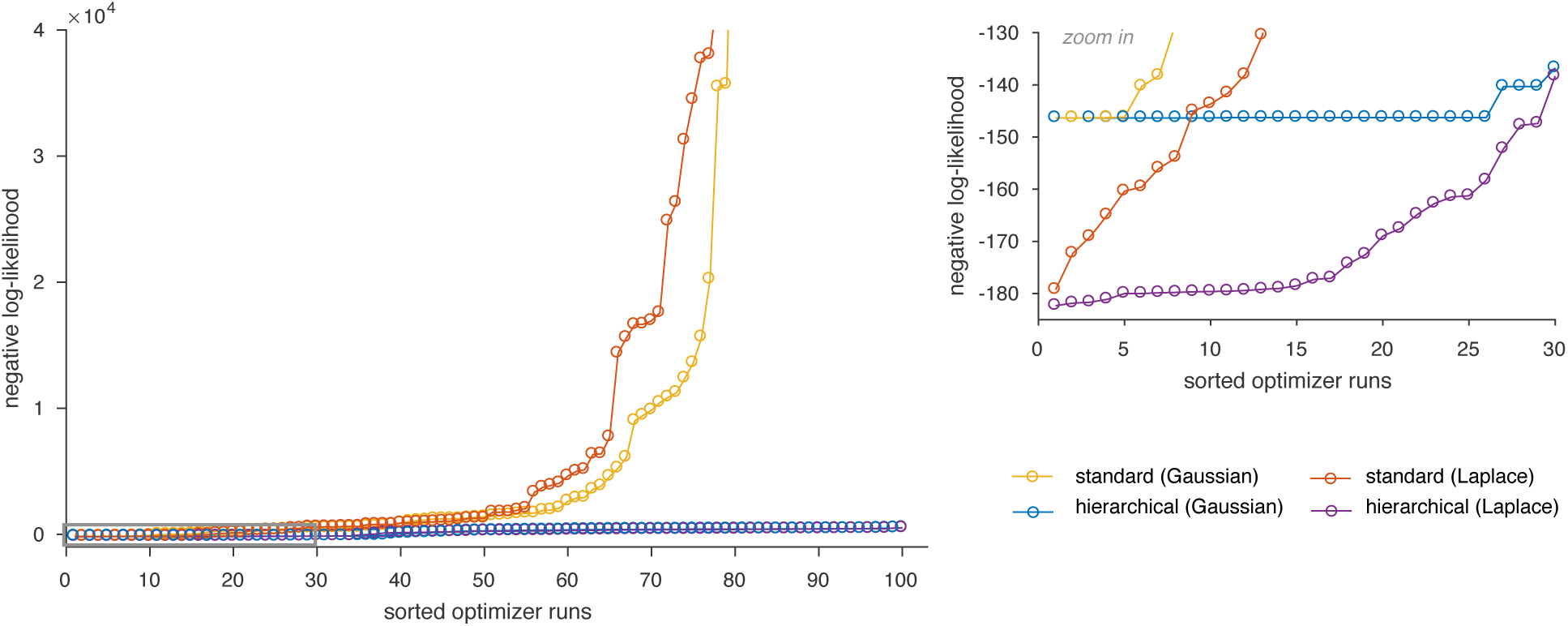
Likelihood waterfall for the JAK-STAT signaling model II.

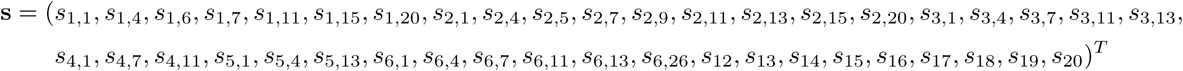

with *n*_*s*_ = 42. The noise parameters do not differ between experiments or replicates, thus, neglecting the subscripts for the experiment-specific condition index *c*_*e*_ and for the replicate index *r*, the noise parameters, which need to be estimated from the data are given by

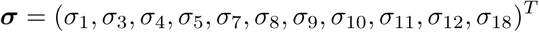

with *n_σ_* = 11. Some observables have the same noise parameters:

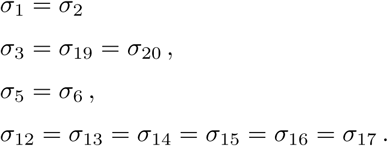

A minor modification from the model proposed by Bachmann et al. (2011) is that the parameterization for the noise of pSTAT5B_au_ did not include an additional parameter for the SOCS3oe experiment, and that the observables for RNA were fitted in linear space. The observable pSTAT5B_rel_ was also fitted on a linear scale, while the other observables were compared at a log_10_ scale (as done by Bachmann et al. (2011)). In our setting, the offset parameters were also multiplied with the scaling parameters, which yielded different optimal values for the offset parameters compared to those found by Bachmann et al. (2011). We performed 100 multi-starts for each optimization approach and noise assumption. The

**Supplementary Table S1:** Overview for the experimental data of JAK-STAT signaling model II.

| name | experiment index $e$ | condition index | condition $\mathbf{u}$ | | | | |
| --- | --- | --- | --- | --- | --- | --- | --- |
| | | | ActD | CISoe | SOCS3oe | SHP1oe | [Epo]/$10^{-6}$ |
| Long | 1 | 1 | 0 | 0 | 0 | 0 | 0.125 |
| Concentration | 2 | 2 | 0 | 0 | 0 | 0 | 0.125 |
| RNA | 3 | 3 | 0 | 0 | 0 | 0 | 0.125 |
| ActD | 4 | 4 | 0 | 0 | 0 | 0 | 0.125 |
|  |  | 5 | 1 | 0 | 0 | 0 | 0.125 |
| Fine | 5 | 6 | 0 | 0 | 0 | 0 | 1.25 |
| CISoe | 6 | 7 | 0 | 0 | 0 | 0 | 0.125 |
|  |  | 8 | 0 | 1 | 0 | 0 | 0.125 |
| CISoe_pEpoR | 7 | 9 | 0 | 0 | 0 | 0 | 0.125 |
|  |  | 10 | 0 | 1 | 0 | 0 | 0.125 |
| SOCS3oe | 8 | 11 | 0 | 0 | 0 | 0 | 0.125 |
|  |  | 12 | 0 | 0 | 1 | 0 | 0.125 |
| SHP1oe | 9 | 13 | 0 | 0 | 0 | 0 | 0.125 |
|  |  | 14 | 0 | 0 | 0 | 1 | 0.125 |
| dose response 7 min | 10 | 15 | 0 | 0 | 0 | 0 | 0.0025 |
|  |  | 16 | 0 | 0 | 0 | 0 | 0.025 |
|  |  | 17 | 0 | 0 | 0 | 0 | 0.25 |
|  |  | 18 | 0 | 0 | 0 | 0 | 2.5 |
|  |  | 19 | 0 | 0 | 0 | 0 | 25 |
| dose response 30 min | 11 | 20 | 0 | 0 | 0 | 0 | 0.0025 |
|  |  | 21 | 0 | 0 | 0 | 0 | 0.025 |
|  |  | 22 | 0 | 0 | 0 | 0 | 0.125 |
|  |  | 23 | 0 | 0 | 0 | 0 | 0.25 |
|  |  | 24 | 0 | 0 | 0 | 0 | 1.25 |
|  |  | 25 | 0 | 0 | 0 | 0 | 2.5 |
| dose response 10 min | 12 | 26 | 0 | 0 | 0 | 0 | 0.0025 |
|  |  | 27 | 0 | 0 | 0 | 0 | 0.0125 |
|  |  | 28 | 0 | 0 | 0 | 0 | 0.025 |
|  |  | 29 | 0 | 0 | 0 | 0 | 0.125 |
|  |  | 30 | 0 | 0 | 0 | 0 | 0.25 |
|  |  | 31 | 0 | 0 | 0 | 0 | 2.5 |
| dose response 90 min | 13 | 32 | 0 | 0 | 0 | 0 | 0.0025 |
|  |  | 33 | 0 | 0 | 0 | 0 | 0.1025 |
|  |  | 34 | 0 | 0 | 0 | 0 | 0.125 |
|  |  | 35 | 0 | 0 | 0 | 0 | 0.25 |
|  |  | 36 | 0 | 0 | 0 | 0 | 2.5 |

parameter boundaries are log_10_(*θ*)_lb_ = −3 and log_10_(*θ*)_ub_ = 3, except for

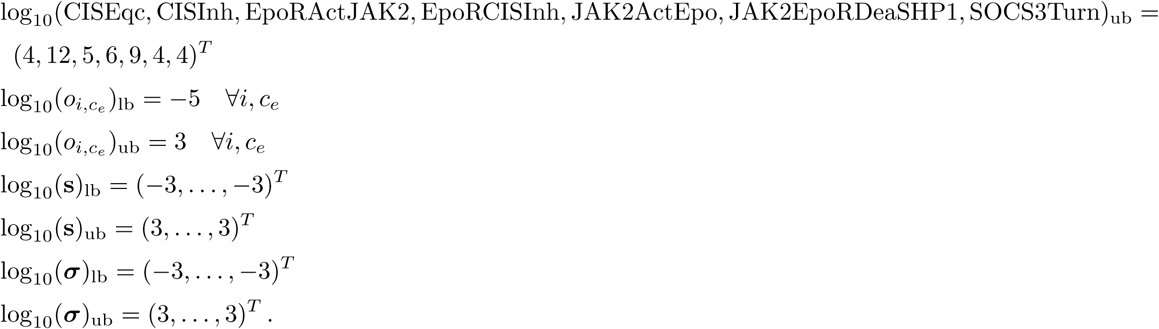

The fitted experimental data for the whole data set are shown in Figure S4. The comparison of Gaussian and Laplace noise showed that Laplace noise yielded a substantially improved fit of the data (Figure S1C).

**Supplementary Figure S4:**
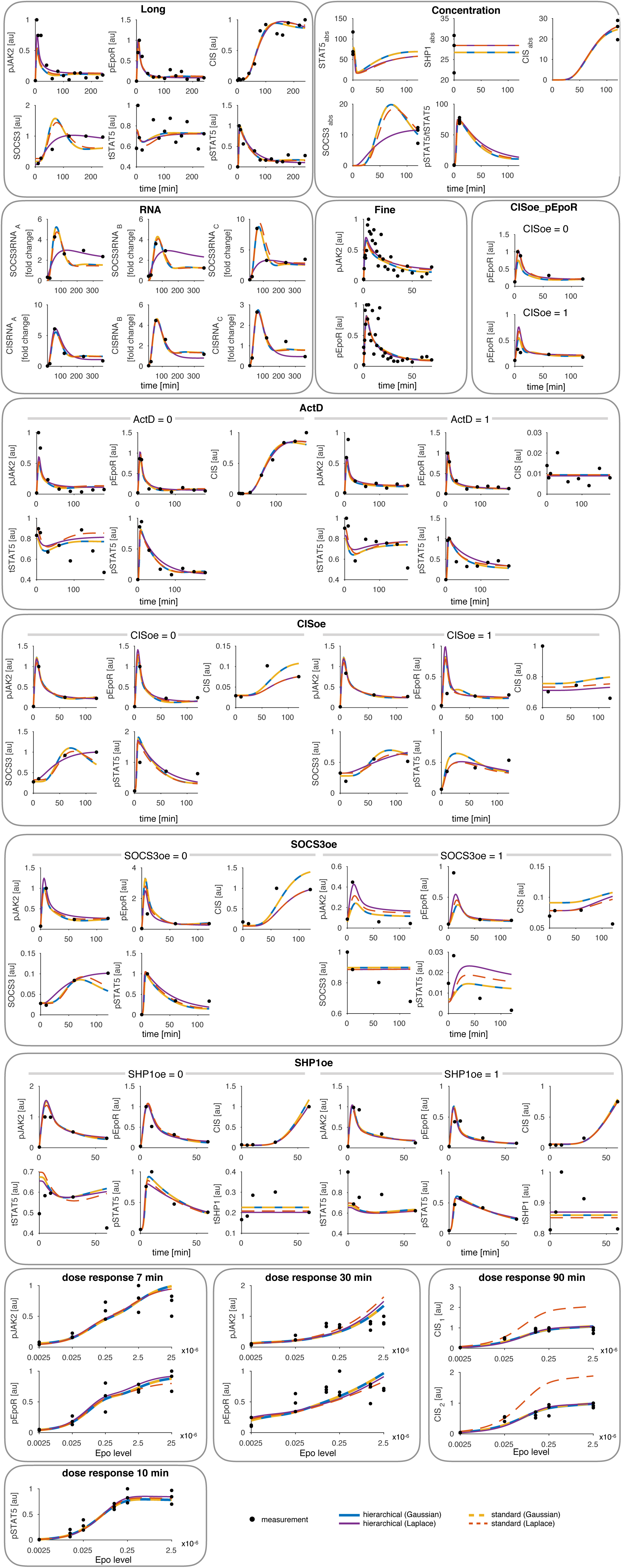
Experimental data for JAK-STAT signaling model II. Boxes indicate different experiments. The lines highlight the different models (Gaussian and Laplace noise) and optimization approaches (standard and hierarchical).

### 4.3 RAF/MEK/ERK signaling

The ODE system for the RAF/MEK/ERK signaling model is given by

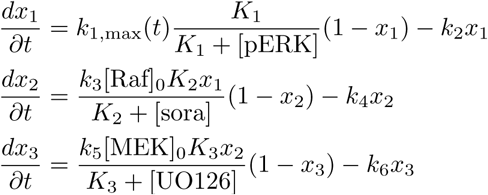

with states *x*_1_ = [pRaf]/[Raf]_0_, *x*_2_ = [pMEK]/[MEK]_0_, and *x*_3_ = [pERK]/[ERK]_0_, and

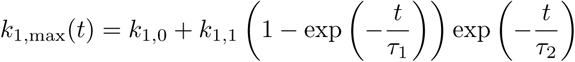

(see (Fiedler et al., 2016) for more details). The initial conditions were assumed to be the steady states reached without stimulation and for *K*_1,max_ = *K*_1,0_. Defining 
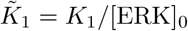

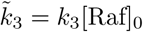
 and

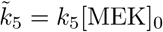
 we obtain

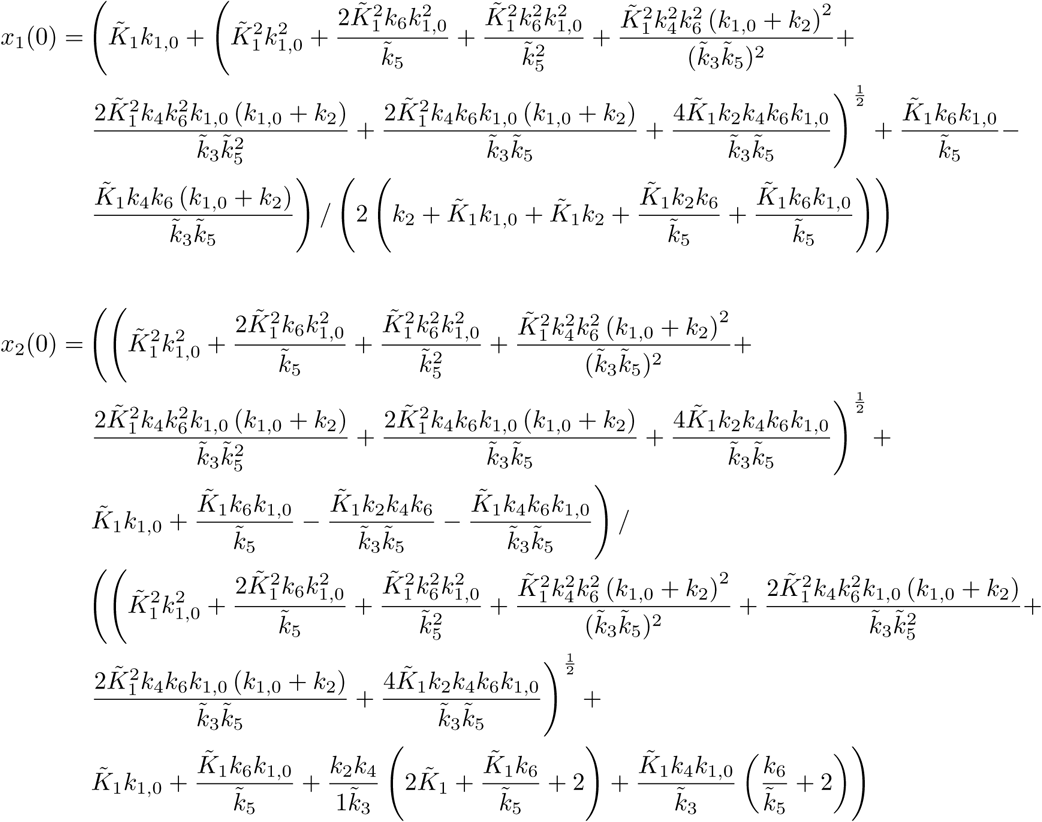

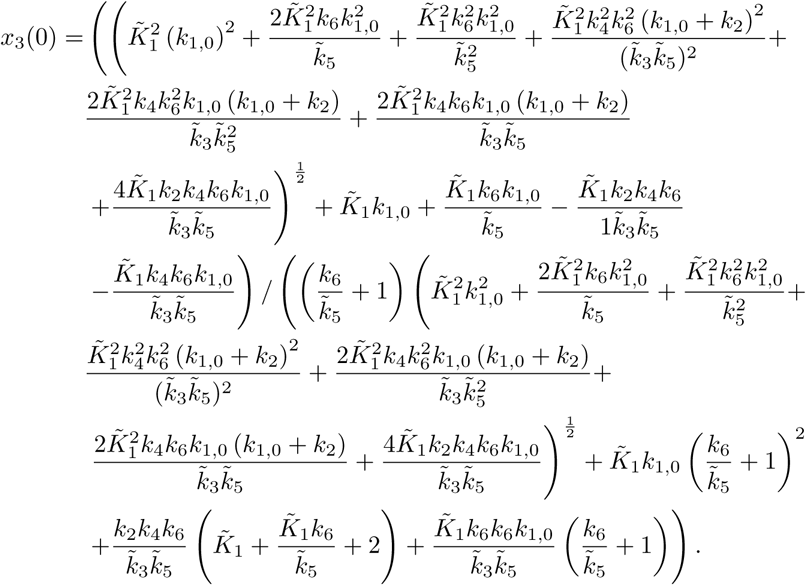

The observables are given by

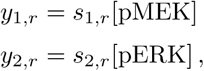

for replicates *r* = 1,…, 4. The indices for conditions and experiments are neglected, since the scaling and noise parameters do not differ for these. The input u describes the concentrations [sora] and [UO126] and the three different conditions are *u*_1_ = (0, 0)^*T*^, *u*_2_ = (0, 30)^*T*^, and *u*_3_ = (5, 0)^*T*^. The parameters, which are estimated from the data, are

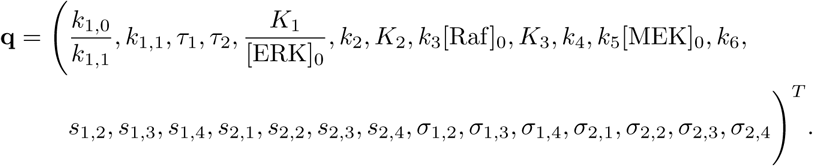

with specific scaling and noise parameters for replicates and observables. The parameters boundaries for the optimization are

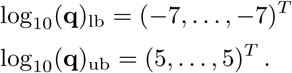

We performed 500 multi-starts to obtain the optimal parameters for both distributions. The comparison between the two noise assumptions for the measurement noise showed that the Laplace noise yielded a substantially better fit (Figure S1C).

**Supplementary Figure S5:**
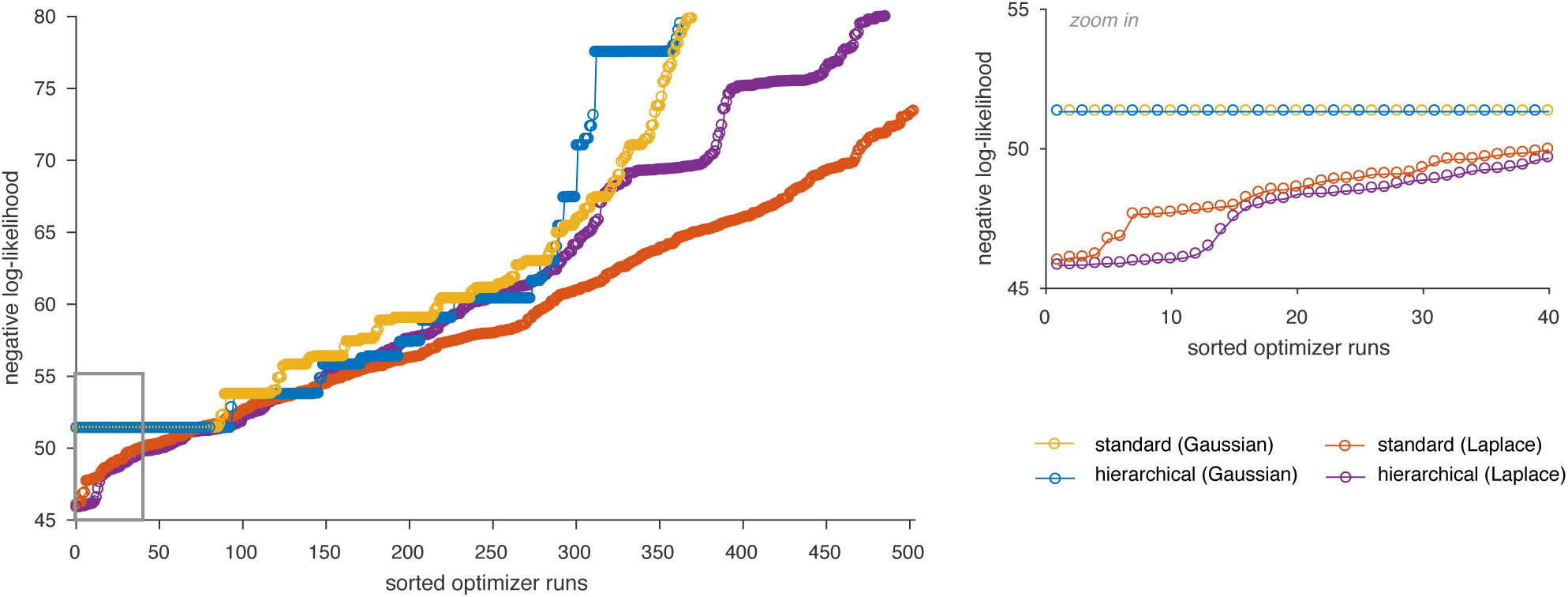
Likelihood waterfall plot for RAF/MEK/ERK signaling.

